# Interaction specificity of clustered protocadherins inferred from sequence covariation and structural analysis

**DOI:** 10.1101/493106

**Authors:** John M. Nicoludis, Anna G. Green, Sanket Walujkar, Elizabeth J. May, Marcos Sotomayor, Debora S. Marks, Rachelle Gaudet

## Abstract

Clustered protocadherins are a large family of paralogous proteins that play important roles in neuronal development. The more than 50 clustered protocadherin isoforms have remarkable homophilic specificity for interactions between cellular surfaces that is controlled by a large antiparallel dimer interface formed by the first four extracellular cadherin (EC) domains. To understand how specificity is achieved between the numerous paralogs, we used a combination of structural and computational approaches. Molecular dynamics simulations revealed that individual EC interactions are weak and go through binding and unbinding events, but together they form a stable complex through polyvalency. Using sequence coevolution, we generated a statistical model of interaction energy for the clustered protocadherin family that measures the contributions of all amino acid pairs in the interface. Our interaction energy model assesses specificity for all possible pairs of isoforms, recapitulating known pairings and predicting the effects of experimental changes in isoform specificity that are consistent with literature results. Our results show that sequence coevolution can be used to understand specificity determinants in a protein family and prioritize interface amino acid substitutions to reprogram specific protein-protein interactions.

## Introduction

Clustered protocadherins (Pcdhs) are a large protein family (53 isoforms in humans) that play roles in vertebrate nervous system development, including neuronal survival, axon targeting, neuronal arborization, and dendritic self-avoidance (1–9). Dendritic self-avoidance is mediated by formation of a clustered Pcdh assembly between two dendrites (10). This assembly relies on individual recognition units formed *in trans* across two cellular membranes that consist of homodimers of the first four extracellular cadherin-repeat (EC) domains in an antiparallel arrangement (**Figure 1A**). These homodimers are highly specific such that no cross interactions are observed in even the most similar isoforms (11–13). The *trans* EC1-4 interaction is also found in other non-clustered Pcdhs (14–16), which have roles in nervous system development and maintenance (17), indicating the importance of this recognition unit in cognitive function. Given that there are many isoforms per vertebrate genome, we sought to understand how specificity is achieved in this large interface.

**Figure 1.**
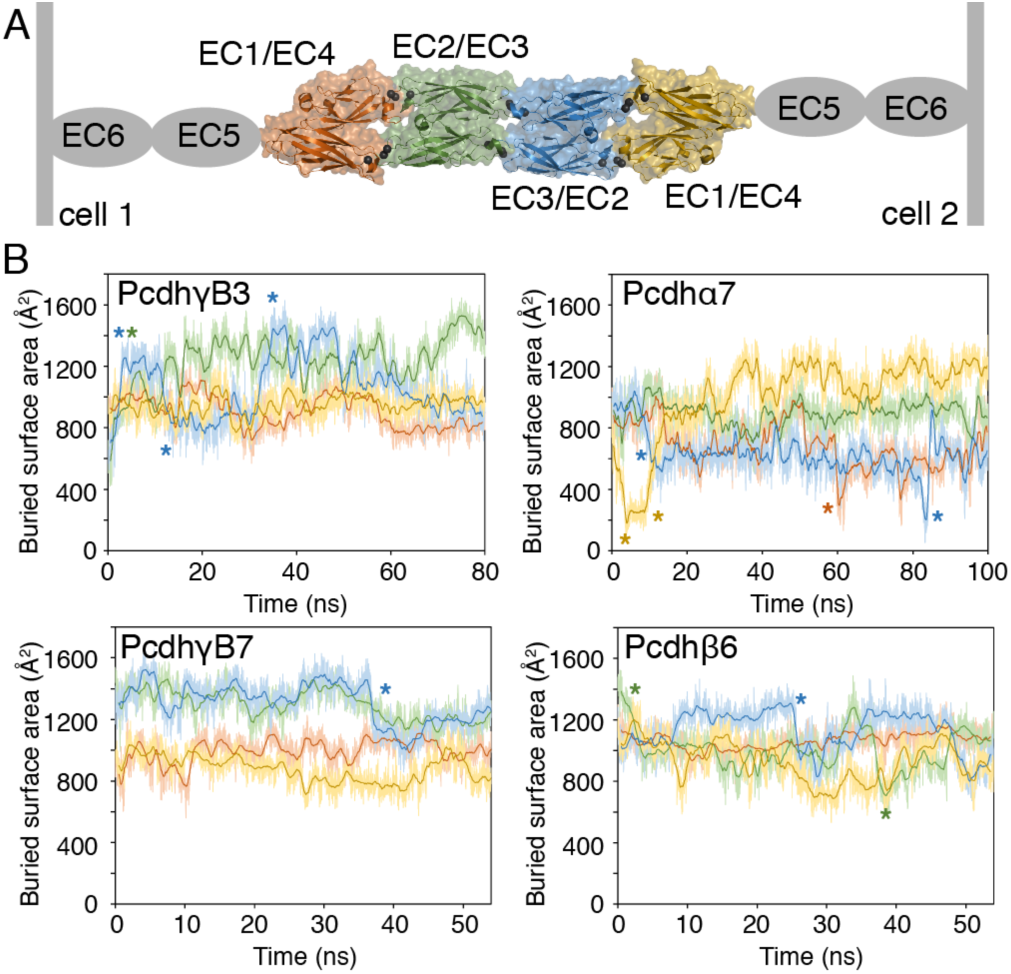
Buried surface area of clustered Pcdh EC interactions fluctuates. (A) The clustered Pcdh dimer is an antiparallel complex of EC1-4 where EC1 of one protomer interacts with EC4 of the other protomer (orange and yellow) and, similarly, EC2 interacts with EC3 (blue and green). Each simulation thus provides two examples of each type of interaction. (B) The BSA of the EC1:EC4 and EC2:EC3 interactions varies throughout the simulations for each of the four simulated isoforms (PcdhγB3 EC1-4, PcdhγB7 EC1-4, Pcdhβ6 EC1-4, and Pcdhα7 EC1-5). Instances where BSA changes rapidly (>400 Å^2^ in under 5 ns) are indicated with an asterisk in the corresponding color.

Structures of these recognition units (14, 18–20) have revealed idiosyncratic characteristics of individual dimer structures, such as the lack of the EC1/EC4 interaction in the structure of PcdhγA1 and PcdhγA8 (20) and the small interface of the EC2/EC3 interaction in PcdhγB3 (14). More subtle structural differences can be observed broadly between the three clustered Pcdh subfamilies, α, β, and γ (14, 19, 20). Based on the variety of interfaces found in the existing crystal structures (14, 18–20), it is possible that every isoform achieves specificity by adopting a different static interface conformation, or that isoforms sample a distribution of conformations, with different combinations of interface residues determining preference for self-interaction. Understanding Pcdh interaction specificity will require disentangling these scenarios by considering both interface conformations and residue-residue interaction preferences.

Prior computational work has sought to understand the evolution of specificity of the Pcdh *trans* interaction, finding positive selection on the *trans* interface (21), and suggesting that the EC2/EC3 interface plays a greater role in specificity between closely related protocadherins (10, 14). However, this computational work did not analyze residue dependencies at the interface, or provide a statistical framework to predict specificities for new mutations or combinations of protocadherins. Recent computational methods based on residue coevolution have proven useful for understanding the structure and function of protein complexes (22, 23). These methods use undirected graphical models of protein sequences to find statistical dependencies between pairs of residues (22, 23), and have succeeded in predicting correct protein-protein interaction pairings for protein families such as bacterial two-component systems (24, 25) and predicting the effect of mutations on protein function (26). Therefore, these generative models of residue dependencies may allow for the prediction of specificities in the Pcdh family.

We use molecular dynamics and new crystal structures to show that isoforms adopt a range of conformations and identify interacting residue pairs. We build a statistical model using evolutionary couplings (22, 23) to analyze specificity of all possible *trans* isoform interactions and infer which domains are important for interaction and specificity. This work provides insight into the molecular origins of specificity within the clustered Pcdh family, and demonstrates that models based on sequence coevolution can be used to guide reprogramming of protein-protein interaction specificity.

## Results

### Molecular dynamics simulations and crystal structures show that Pcdh *trans* interfaces sample a distribution of conformations

The *trans* interface of clustered Pcdhs is specific for self-interaction (11, 12). Crystal structures of many Pcdh isoforms dimers (14, 18–20) revealed idiosyncratic features of individual isoforms, raising the possibility that different isoforms adopt different conformations *in vivo*, resulting in molecular specificity. For example, PcdhγA1 (in one of two dimers) and PcdhγA8 (20) lack EC1/EC4 contacts, and PcdhγB3 EC2–3 has a surprisingly small interface (14). We thus used MD simulations to test whether the distinctive features of individual isoforms were the result of static conformational differences, or the result of a dynamic interface crystallized in different conformations.

We performed all-atom equilibrium simulations of four different EC1-4 homodimers from the different Pcdh subfamilies to distinguish the above hypotheses. We simulated clustered Pcdh isoforms PcdhγB3, PcdhγB7, Pcdhβ6, and Pcdhα7 (**Table S1**). Overall, the complexes did not dissociate over the course of simulation (**Figure S1A**, **Figure 1A**). Between 5 and 54 ns, the average BSA was 4600 ± 300 Å^2^ for PcdhγB3, 5100 ± 200 Å^2^ for PcdhγB7, 4300 ± 200 Å^2^ for Pcdhβ6, and 3400 ± 300 Å^2^ for Pcdhα7. Between individual EC domain dimers, which average 700-1300 Å^2^ in BSA, fluctuations in BSA can be more than 400 Å^2^ over the course of a few nanoseconds (**Figure 1B**) (**Table S2**). For example, one EC1/EC4 dimer of Pcdhα7 fluctuates in BSA from 1000 Å^2^ at the start of the simulation to ~200 Å^2^ at 6 ns then to 1200 Å^2^ around 35 ns (**Figure S1**).

The published structure of PcdhγB3 has a particularly small BSA at the interface (12). The fluctuation in EC2/EC3 BSA during simulation indicates that perhaps the EC2/EC3 interface was occluded by HEPES during crystallization (14, 20). We determined new structures of PcdhγB3 EC1-4 both in Tris buffer and in a lower concentration of HEPES (50 mM vs. 100 mM) (**Table S3**). These two structures are nearly identical to the original (overall RMSD (over 3210 atoms): 0.890 and 0.785 Å, respectively) (**Figure 2A**). The corresponding electron density maps show less or no density in the location of the HEPES molecule in the original structure (**Figure 2B**). Thus, HEPES was not the cause of the low BSA of the EC2/EC3 interface, but rather, this structure represents an accessible conformation of this Pcdh dimer.

**Figure 2.**
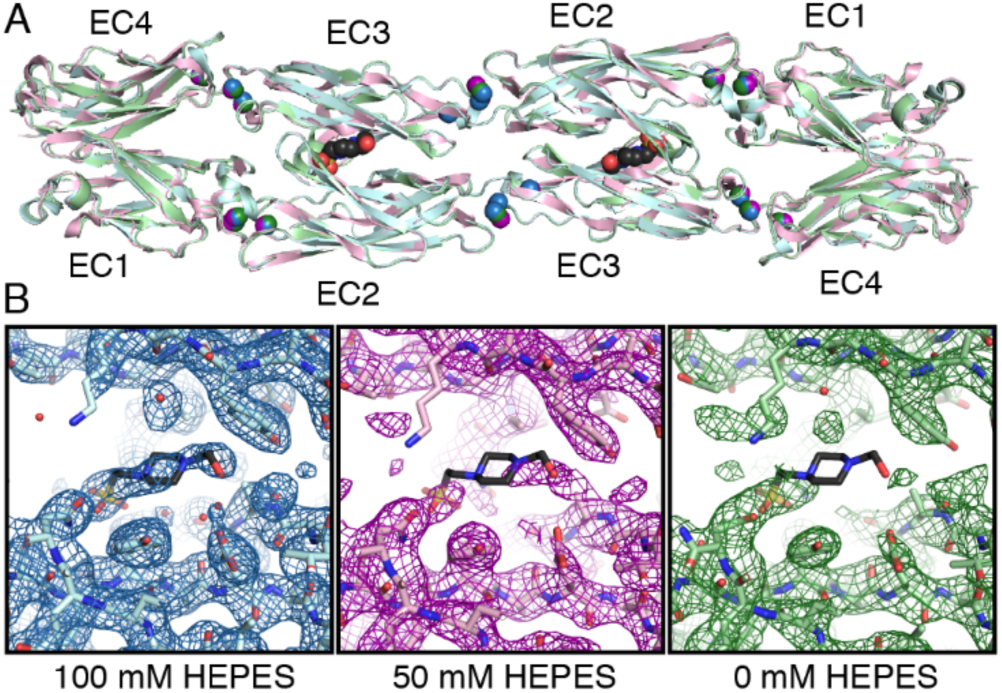
HEPES does not influence the structure PcdhγB3 EC1-4. (A) Crystal structures of PcdhγB3 EC1-4 in the presence of 100 mM HEPES (blue; PDB ID: 5k8r), 50 mM HEPES (pink; PDB ID: 6mer), and no HEPES (green; PDB ID: 6meq) adopt the same conformation. The HEPES molecule in the original structure (PDB ID: 5k8r) is shown in black and red. (B) Composite omit maps (which reduce model bias in electron density) of these three structures show density for HEPES in the original structure, but less or no density in the lower and no HEPES conditions.

The increase in the EC2/EC3 interface BSA in the PcdhγB3 EC1-4 simulation is predominantly due to a conformational change of the β4-β5 loop of EC2, which moves closer to EC3 of the other protomer in the first 6 ns of the simulation (**Figure 3A**). The conformational change leads to the formation of hydrogen bonds between Y161 and K302, and N155 and T286, and a van der Waals interaction between L156 and M216 (**Figure 3B**). The corresponding inter-residue distances vary in a coordinated fashion as the loop changes conformation between the disengaged state seen in the crystal structure and an engaged state where the loop interacts directly with EC3 (**Figure 3C**). The structural heterogeneity of the β4-β5 loop of EC2 is echoed in the diverse conformations of this loop in other clustered Pcdh structures (**Figure S2**).

**Figure 3.**
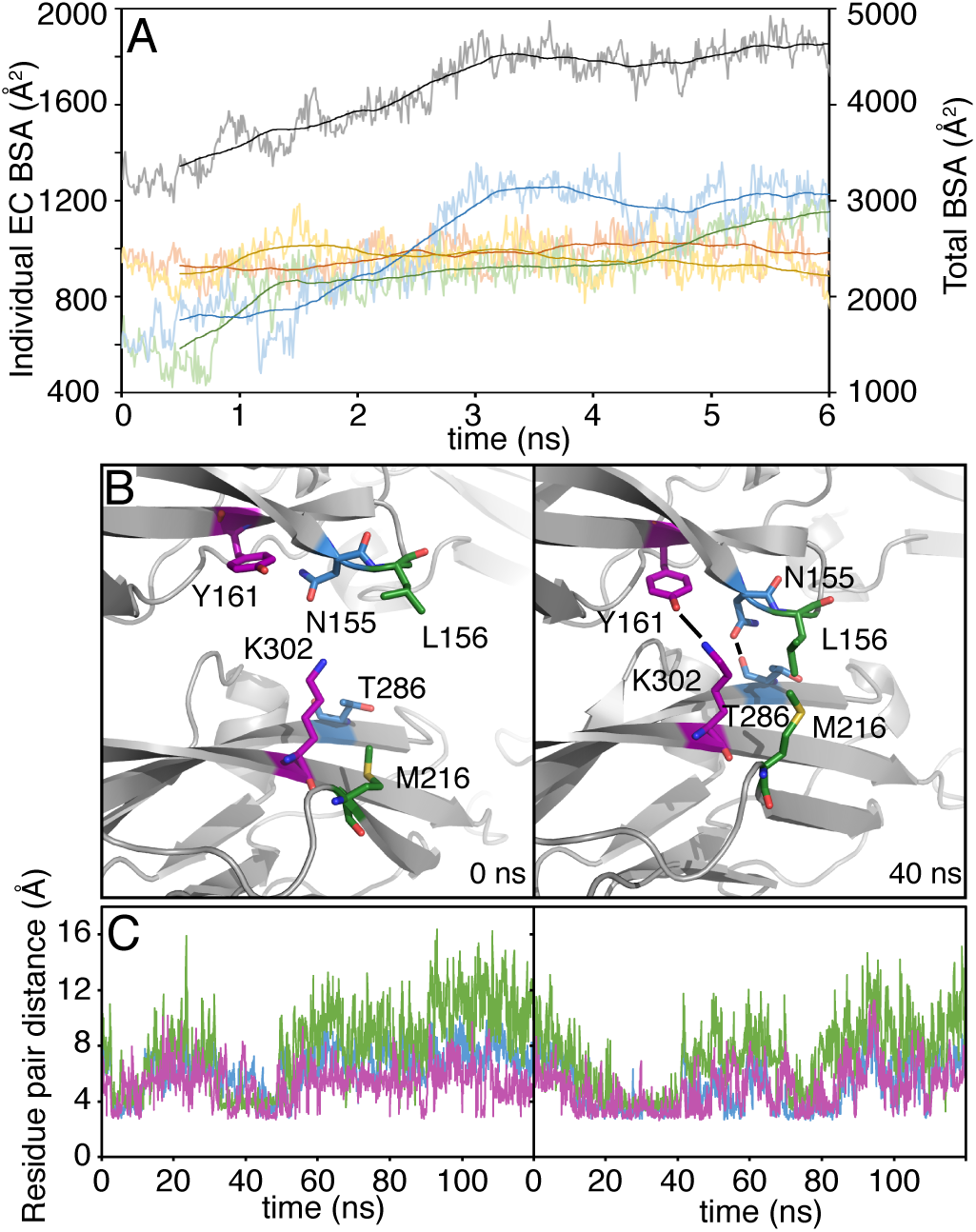
PcdhγB3 EC1-4 EC2:EC3 rearranges to interact extensively in molecular dynamics simulations. (A) BSA values plotted for the first 6 ns show that the overall BSA of the dimer increases from ~3000 Å^2^ to 4500 Å^2^ in the first 3 ns of the simulation (black). This increase in BSA is due to independent increases at the EC2:EC3 interfaces (blue and green), while the EC1:EC4 interfaces maintain the same BSA (yellow and red). (B) In particular, the EC2 β4-β5 loop, containing N155, L156 and Y161, changes conformation from 0 (left) to 40 ns (right) to interact with T286, M216 and K302, respectively, in EC3 of the other protomer. (C) Throughout the simulation, these residue pairs (N155:T286, blue; L156:M216, green; Y161:K302, magenta) fluctuate between an engaged conformation with residue-residue distances (calculated as the two closest non-hydrogen atoms) at 3-4 Å, and a disengaged state with much larger residue-residue distances. The two plots represent the two instances of EC2:EC3 interactions in the homodimer.

The fluctuations seen in MD suggest that if these individual EC interactions were found in isolation they would be low affinity, consistent with the observation that constructs of EC1-3 do not dimerize in solution (13, 18). These fluctuations in BSA agree with other simulations that find sharp decreases in BSA as prerequisites to protein complex dissociation (27). Together, these observations indicate that the four individual EC domain dimers provide weak affinity to the complex and that the polyvalency of the interactions increases the avidity. Polyvalent interactions are known to increase both the strength and specificity of molecular recognition (28). Overall, our simulations indicate that each isoform can sample a range of interface conformations, and that the available crystal structures represent only a snapshot of these conformational possibilities.

### Highly coevolving residue pairs are frequently in contact in simulations

Computational methods based on residue coevolution predicted the EC1/EC4 interaction and established that the *trans* dimer architecture would be found in non-clustered Pcdhs, both findings later confirmed experimentally (14, 15, 18, 29). Sequence coevolution methods for protein-protein interface determination are typically benchmarked by comparing highly coevolving residue pairs that are not due to intramolecular structural features to their inter-residue distances in experimentally-determined structures (30, 31). However, coevolving residues can correspond to positions only in contact in certain conformations (32, 33), opening up the possibility that intermolecular coevolving residue pairs may also only be present in some conformation. Given the dynamic nature of the Pcdh interface and the above observation that crystal structures represent only a snapshot of possible conformations, we analyzed how often highly coevolving interface residues in the Pcdhs (14, 18) are in contact across simulations and over time.

In our sequence coevolution analysis, the set of top 200 coevolving pairs includes mostly intramolecular contacts and 15 intermolecular pairs (**Figure S3**). We calculated the residue-residue distance of these 15 coevolving intermolecular pairs over the course of the MD simulations (**Figure 4**). Of note, each simulation provides two (semi-)independent observations for each residue pair due to the two-fold symmetric nature of the dimer. In general, residue pairs with higher coevolution scores are more frequently in contact in more of the simulations, consistent with the observation that lower-scoring residue pairs are less predictive of physically interacting residues (30, 31).

**Figure 4.**
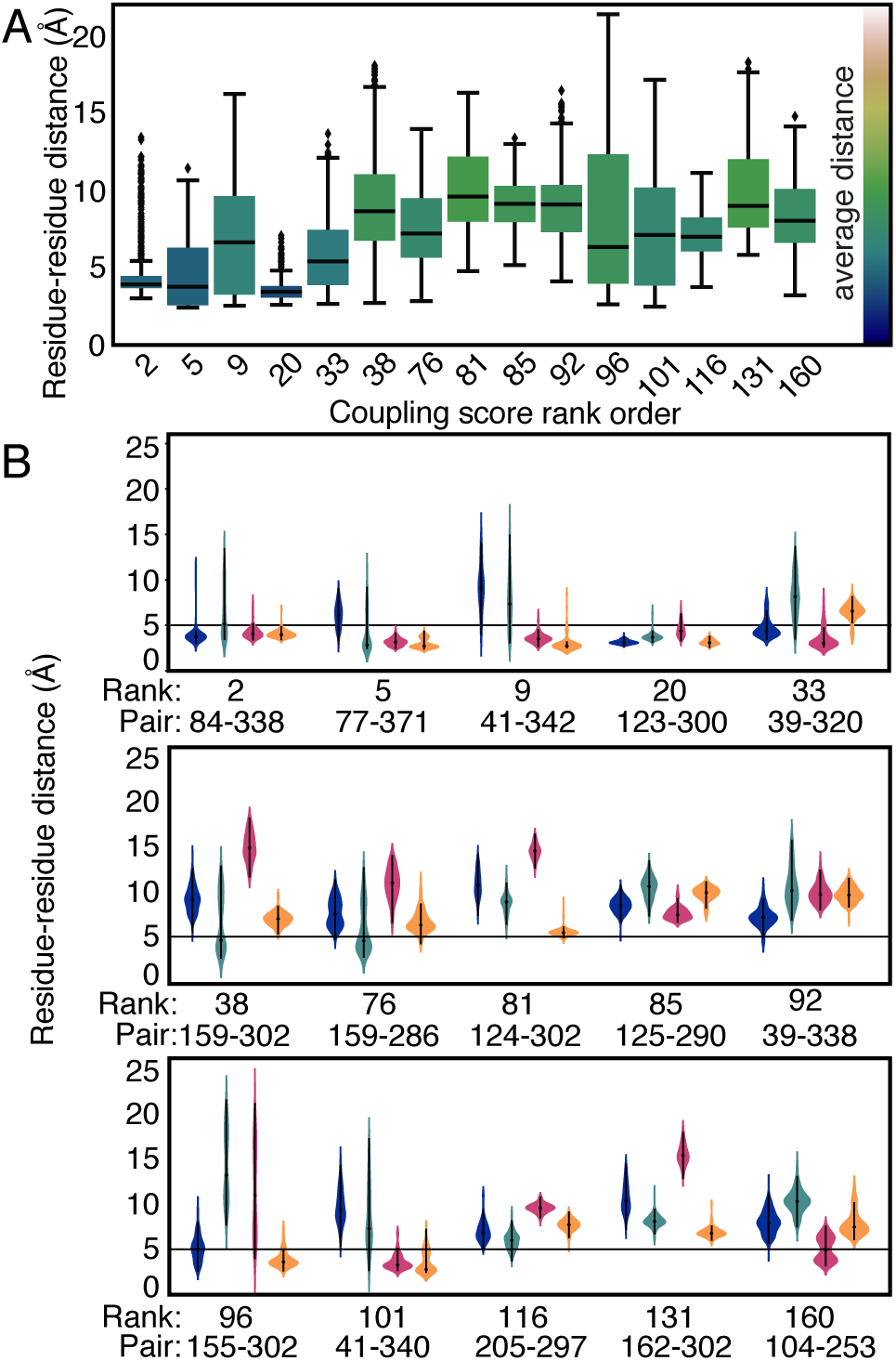
Evolutionary coupling scores of intermolecular pairs correlate with residue-residue interactions in simulations. (A) Box plots of the residue-residue distances of top 15 intermolecular pairs for all four simulations. Boxes are shaded based on the mean residue-residue distance (scale on right). (B) Violin plots of residue-residue distances of the top 15 intermolecular pairs (listed by rank order and pair positions) for the PcdhγB3 EC1-4 (dark blue), PcdhγB7 EC1-4 (orange), Pcdhβ6 EC1-4 (magenta), and Pcdhα7 EC1-5 (teal) simulations. Distances were sampled at 1 ns and both intermolecular pairs were included in each distribution (e.g. 84-338’ and 84’-338).

A closer examination of the top 15 intermolecular pairs reveals diverse trajectories during simulations. For the most highly coevolving pairs, the residues are in close contact for most of the time in most of the simulations. For example, the 84-338 and 123-300 pairs (based on PcdhγB3 numbering in **Figure S4**) remain at ~4 Å throughout all but one simulation (**Figure S5**). For some of the lower-scoring residue pairs, the residues are in close proximity in some of the simulations but further in others, e.g. the 159–302 pair has residue-residue distances that fluctuate between 6 and 18 Å in most simulations but stay consistently close in the Pcdhα7 interface. Other lower scoring pairs fluctuate widely in all simulations and rarely if ever come into contact, such as the 39-338 pair.

Residue coevolution analysis thus captures residues that are frequently in contact in different protocadherin isoforms and during sampling of different conformations. Strongly coevolving residue pairs are more likely to interact consistently over time but different homologous structures do not exhibit all of these interactions. Consequently, some coevolving pairs could be missed from analyzing isolated crystal structures of single isoforms. These results indicate that sequence covariation identifies residues that are important to the Pcdhs *trans* dimer interface, and that these residues can be used to predict residue pair contributions to interaction specificity.

### Statistical energy of interaction describes Pcdh specificity distributions

We used evolutionary couplings to build a model of clustered Pcdh interaction specificity (see Supplementary Information). Evolutionary couplings are calculated using an undirected graphical model, which has parameters for single-site biases and pairwise residue preferences for all sites (22, 23). These parameters have been used successfully to infer the effects of mutations and predict correct pairings for other paralogous interacting proteins (24–26). Here, we assess the propensity for any two Pcdhs to interact by summing the pairwise residue preferences for all interface residue pairs, producing a score which we call the statistical energy of interaction (SE*_i_*, see Methods and **Figure 5A**). A higher SE*_i_* indicates a higher propensity for interaction.

**Figure 5.**
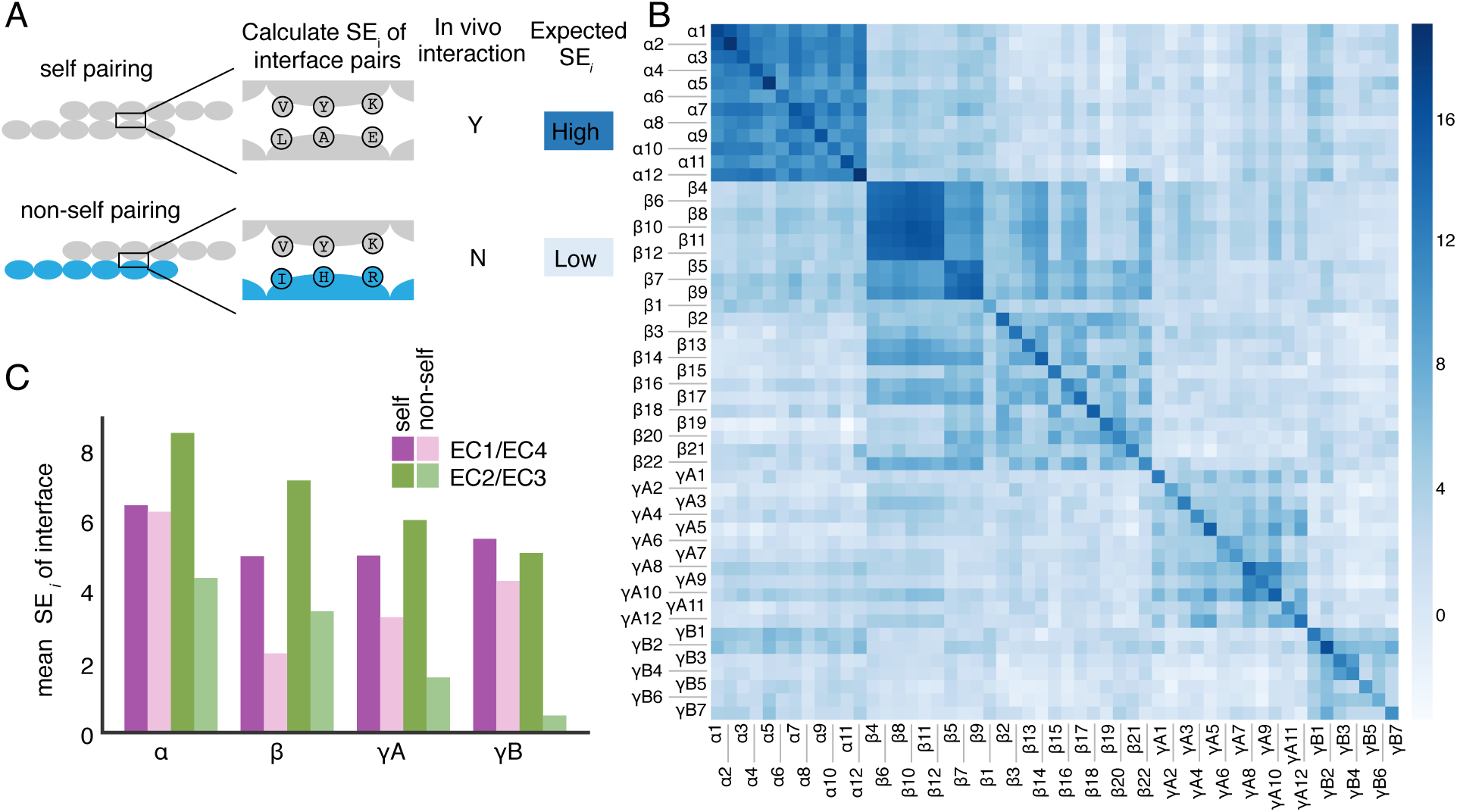
Statistical energy of interaction supports experimental evidence for self-interaction and divergent roles of EC2/EC3 and EC1/EC4 interfaces. (A) Statistical energy of interaction (SE*_i_*) for every possible pairing of Pcdhs was calculated by summing the energy contribution of each interacting residue pair at the interface. Based on prior work (24, 25) we expect to observe high statistical energy for self-pairings, which interact *in vivo* (11, 12) and low statistical energy for non-self pairings. (B) Statistical energy for all possible combinations of isoforms. (C) Mean statistical energy of every self and non-self pairing for the EC2/EC3 and EC1/EC4 interface, measured within all subfamilies of clustered Pcdhs in mouse.

We observe that for the α, β, and γ Pcdh subfamilies in mouse, the SE*_i_* of a sequence with itself (a self pairing) is higher than the SE*_i_* of a sequence with a different isoform (a non-self pairing; **Figure 5B**). This is generally consistent with previous cell aggregation experiments in which clustered Pcdhs only form homodimers (11, 12). While these studies observed no non-self interaction in their experimental systems, our model finds that in some cases the SE*_i_* for non-self pairs is as high as for a self pairing, e.g. between the β4 and β6 isoforms. This could be due to particulars of the cell aggregation assay, or suggest that some *in vivo* determinants of specificity are not fully captured by our model.

The SE*_i_* computed using evolutionary couplings confirms our earlier finding (14) that EC2/EC3 interface tends to contribute more to specificity than the EC1/EC4 interface. The non-self Pcdh pairs of the mouse α, γB, γA families have a lower SE_i_ in the EC2/EC3 interface than they do in the EC1/EC4 interface, indicating that the EC2/EC3 interface contributes more to specificity of isoforms (**Figure 5C**, **Figure S6**). The α subfamily has nearly identical SE*_i_* between EC1/EC4 self and non-self interfaces. The difference in SE*_i_* between EC2/EC3 and EC1/EC4 non-self pairs may be due to a greater number of mutations between self and non-self EC2/EC3 pairs compared to EC1/EC4 pairs, which negatively correlates with the SE*_i_* (**Figure S7**).

Our model parameters are inferred only from natural sequences, and therefore self-pairings, which may bias the model against non-self pairings. In order to avoid this possible bias, we set up an iterative pairing algorithm which allows the isoforms to find favorable non-self pairings, if such pairings exist (23, 24). First, isoforms are paired randomly within the same species to create an initial sequence alignment that eliminates the self-pairing bias. Then, each iteration of the algorithm entails inferring the parameters of the evolutionary couplings model from the current alignment, and then updating the isoform pairing in the alignment according to which pairs have the highest SE*_i_* (see Supplemental Information, **Figure S8**). The algorithm reproduces self pairings for 74% of all sequences in the alignment, averaged across five replicates, after iteration to convergence. This is on par with accuracy of partner detection with other proteins pairs performed by related algorithms (23,34), and supports use of our model trained on natural sequences.

We then used the iterative pairing algorithm to test our observation that the EC2/EC3 interface is the main determinant of specificity. When sequences were paired based on the SE*_i_* from only their EC1/EC4 or EC2/EC3 interfaces, we find 49% and 59% accuracy on all sequences in the alignment, respectively (**Figure S8**). This confirms that the EC2/EC3 interface has more discriminatory power between isoforms, but that both interfaces act in combination to achieve full specificity of the interface.

Our statistical energy of interaction model allows us to compute a statistical energy of interaction score for all pairs of clustered protocadherins which generally agrees with experimental findings about specificity of protocadherin isoforms. Importantly, it also allowed us to dissect contributions of various interface components at an overall and subfamily level. We observed lower mean SE*_i_* at the EC2/EC3 non-self interface than at the EC1/EC4 non-self interface and better pairing of EC2/EC3 interactions than EC1/EC4 interactions in our matching algorithm, indicating that the EC2/EC3 interface tends to be more involved in specificity.

### Statistical model explains changes in Pcdh specificity in chimera mutants

The statistical energy of interaction model allows us to predict how particular mutations may alter interaction specificity by recalculating SE*_i_* using J*_ij_* terms from the mutant sequence. Previous work has tested chimeric constructs in cell aggregation assays to understand how specificity is encoded in the clustered Pcdh family (13). In these experiments, chimeric constructs were tested for interaction with one of the parent isoform (which we will call the ‘target’ isoform). The original chimeras did not interact with the parent but were mutated such that they could form an interaction with the target isoform. To determine how these mutations affect the specificity we calculated the SE*_i_* between the chimera and target isoforms, and the mutant chimera and target isoforms (**Figure 6A**). If the change in SE*_i_* from the chimera to the mutant chimera (ΔSE*_i_*) is positive, it means the mutant chimera is more likely to interact with the target isoform than the original chimera. If ΔSE*_i_* is negative, the mutant chimera would be less likely to interact with the target isoform than the original chimera. In Rubinstein et al. (2015), three pairs of closely related isoforms (>85% identity) were chosen: Pcdhα7/Pcdhα8, PcdhγA8/PcdhγA9 and Pcdhβ6/ Pcdhβ8 (13). Out of the seven mutant chimeras that were able to interact with the target isoform, we correctly predicted five out of seven, observing that the introduced mutations result in an increase in SE*_i_* (**Figure 6B**). Of the two we did not predict, one is a single site mutation in EC1 in a Pcdhβ6/Pcdhβ8 chimera that had almost no change in calculated SE*_i_*. The other exception was a set of mutations in the EC3 β6-β7 loop converting the PcdhγA8 to the PcdhγA9 sequence, which we predicted to decrease SE*_i_*. This region has many insertions and deletions, creating gaps in the multiple sequence alignment. These gaps are treated as missing data by our statistical energy model and therefore reduce the quality of the predictions. Overall our statistical energy model of interaction specificity, when enough high-quality sequence information is available, is very consistent with available experimental data.

**Figure 6.**
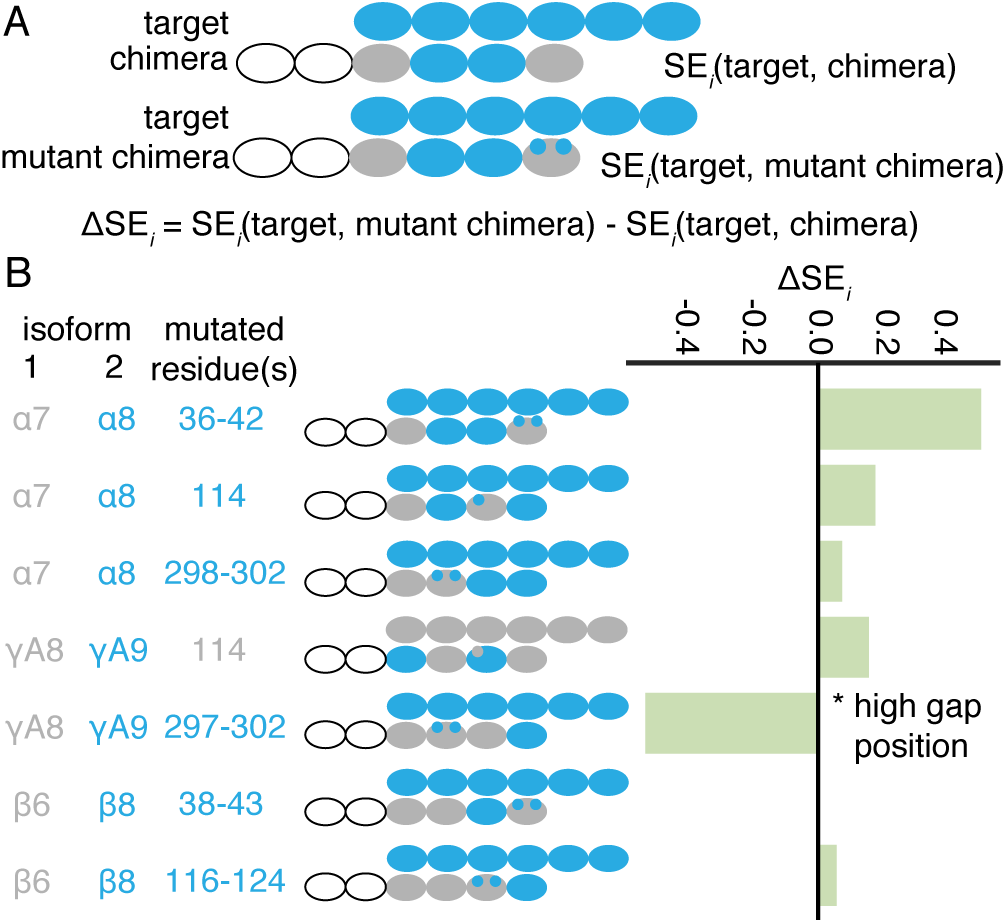
Statistical energy of interaction is corroborated by changes in specificity in chimera mutants. (A) Schematic of calculation of statistical energy difference between chimera sequences and mutant chimera sequences. (B) Chimeras from (13) and our computed statistical energy for each pairing. Mutated residues are numbered according to Figure 5 of (13). The γA8/γA9 297-302 pair was not predicted well due to gaps in the alignment in this residue range.

## Discussion

The highly specific antiparallel Pcdh interface that forms between neurons is required for many roles in neuronal development. The results presented here address the determinants of specificity in this interaction by simulating the dynamics of the Pcdh interface and modeling the contributions of each residue pair to interaction specificity.

Our MD simulations of clustered Pcdh dimers reveal that individual EC interactions sample a range of conformations in every isoform. These simulations, combined with crystal structures, show that the surprisingly small EC2/EC3 interface of our previous structure of a PcdhγB3 EC1-4 homodimer was not due to the presence of HEPES as previous suggested (20), but rather the conformation seen in the crystal structure is part of the normal dynamics of the conformational landscape. The variations in BSA for individual EC interactions suggest that the individual interactions are weak, and the overall stability of a Pcdh dimer is established by the polyvalent nature of these individual EC interactions. This type of cooperative binding is widespread in biology and plays roles in multi-subunit protein machine assembly, signaling at the membrane, and signaling between cells (28, 34, 35).

The MD simulations allowed us to observe how evolutionarily coupled residue pairs across a protein interface vary over time. Overall, we found that higher scoring pairs are closer together throughout the simulations and across multiple homologs, concurring with previous empirical results that coupled pairs are more likely to be close in 3D (30, 31), and that evolutionary couplings can correspond to multiple incompatible conformations (32, 33). Residue-residue distances for these evolutionarily coupled pairs can vary greatly over time, indicating that static structures in single conformations may miss direct interactions of coupled pairs. This knowledge could inform further developments for benchmarking structure prediction using coevolution data.

We constructed a model of interaction specificity from sequence data, using residue pairs found to interact in our simulations. We used this model to evaluate pairs of individual EC interfaces and found that the statistical energy of interaction (ΔSE*_i_*) supports literature results that the Pcdh interface is specific for self-interaction. We find that non-self pairings of the EC2/EC3 interface have lower SE*_i_* than non-self EC1/EC4 pairings for the α, γA and γB subfamilies, indicating that the EC2/EC3 interface has a greater contribution to specificity. There are some differences between subfamilies as noted previously (14, 20), with the β and γA subfamilies having nearly equal contributions to specificity from both interfaces. Our iterative pairing algorithm corroborates this result, finding that the EC2/EC3 interface is more discriminatory than the EC1/EC4 interface.

We validated our model by predicting the effects of mutations in chimeric constructs that changed specificity. These results establish that sequence coevolution analysis can be used to prioritize specific sites/pairs in order to reprogram protein-protein interactions.

The work presented here suggests a strategy used by clustered Pcdhs to ensure specificity and yet allow new specificities to easily arise through evolution. Small changes in individual EC affinity caused by a small number of mutations can alter the affinity of the whole dimer through the cooperativity of the individual EC interactions. This strategy may explain the pervasiveness of this interface for cell-cell adhesion in nervous system development.

## Methods

Detailed procedures for all methods described here are provided in the **SI Appendix**.

### Construction of a statistical interaction energy model of clustered Pcdh specificity

We used evolutionary couplings to build a model of clustered Pcdh interactions. Previous studies used the statistical energy of an evolutionary couplings model to identify interacting histidine kinase-response regulator pairs (24, 25) and to predict the effects of mutations on protein function (26). For our model, only the interface residue pairs determined by our molecular dynamics approach were used. The interaction energy between two sequences (σ^A^, σ^B^) is the sum of the individual coupling terms (*J_ij_*) between the interface residues of the two sequences:

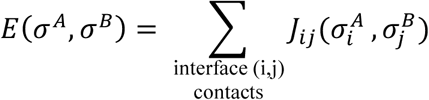

The *J_ij_* term is the matrix of pairwise residue preferences for all possible amino acids in positions i and j (26). The change in *E(σ)* is used to predict whether the interaction will become more or less favorable. See **SI Datasets** for the interface residues, the residue pairs, and an alignments of mouse isoforms. Model parameters are available on request.

To determine the single mutations most likely to reprogram clustered Pcdh A to interact with clustered Pcdh B, we first calculated the statistical interaction energy of A with A and of A with B. We then computationally swap in each residue, one at a time, from B into A, and assess interaction energy of this new A* chimera with sequence A and with sequence B. We compute a change in energy (ΔE) between this mutant sequence and the wild type interaction energies. We excluded the C-type isoforms from this analysis because they are evolutionarily distinct and have unique biological functions (36–38).

## Supporting information

## Acknowledgements

We thank members of the Gaudet and Marks labs for stimulating discussions on this project. We thank Raul Araya-Secchi and Bennett Vogt for initial work on PcdhγB3. We thank the beamline staff at NE-CAT and the support team at SBGrid for help with crystallographic data collection and analysis. NE-CAT is funded by NIH (P41 GM103403 and S10 RR029205) and the Advanced Photon Source by the US Department of Energy (DE-AC02-06CH11357). Evolutionary couplings analysis was conducted on the Orchestra High Performance Compute Cluster at Harvard Medical School, which is funded by the NIH (NCRR 1S10RR028832-01). All MD simulations were performed on TACC’s Stampede2 supercomputer (XSEDE MCB140226) and OSC’s Owens cluster (PAS1037). JMN was supported by a National Defense Science and Engineering Graduate Fellowship, AAG by National Science Foundation Graduate Research Fellowship DGE1144152. Support was also provided by the National Institutes of Health (GM106303 to DSM; DC015271 to MS).

