## Supplementary material for "Interaction specificity of clustered protocadherins inferred from sequence covariation and structural analysis"

**This PDF file includes:**

Supplementary text
Figs. S1 to S9
Tables S1 to S4
References for SI reference citations

**Other supplementary materials for this manuscript include the following:**

Datasets S1 to S3

### Supplementary Information Text

### Experimental Methods

**Crystallography of PcdhγB3 EC1-4 without HEPES**. PcdhγB3 EC1-4 was purified as previously reported (1) and crystallized in 10% PEG 5000 monomethylester and 4% ethylene glycol with either 50 mM HEPES pH 7 or 100 mM Tris pH 7. Crystals were cryo-protected in reservoir plus 20% glycerol and cryocooled in liquid $N_2$. X-ray data were collected according to **Table S3** and processed in HKL2000 (2). Structures were determined using molecular replacement of the published PcdhγB3 EC1-4 structure (1). Model building was done in COOT (3). Refinement and generation of composite omit maps was done in PHENIX (4). Software is maintained through SBGrid (5). Refinement and model statistics are listed in **Table S1.**

**MD simulations**. Four different crystal structures of clustered Pcdhs – mouse Pcdhα7 (5dzv), β6 (5dzx), γB7 (5szp) and human PcdhγB3 (5k8r) were each solvated using TIP3P water (Table S3). All the systems were neutralized and ionized with 150 mM NaCl. Resulting systems were minimized for 5,000 steps and equilibrated for 200 ps with backbone constraints ($k$ = 1 kcal/mol/) and for an additional 1 ns without these constraints. During these initial 1.2 ns of simulation a Langevin damping coefficient of $\gamma = 1.0$ ps$^{-1}$ was used. Subsequent dynamics used $\gamma = 0.1$ ps$^{-1}$. All these simulations were performed using NAMD 2.12, the CHARMM36 force field (6, 7), and the $NpT$ ensemble with $T$ = 300 K and $p$ = 1 atm.

Buried surface area (BSA) was calculated every 10 ps for each trajectory using the “measure sasa” tool in VMD (8) with 1.4 Å sampling radius. We first computed the solvent accessible surface area (SASA) of individual protomers, and then the SASA of the complex formed by that pair. The final BSA reported was calculated by subtracting the SASA of the complex from the addition of the SASAs of each individual subunit.

**Defining interface residues using structural analysis and molecular dynamics simulations**. To build a model for Pcdh interaction specificity, we needed to define a set of residues that constitute the interface for clustered Pcdhs because using the full covariation matrix for all residues would be computationally prohibitive. First, we determined the interface residues (defined as closest heavy atoms are within 8 Å) from all available crystal structures of validated antiparallel clustered Pcdh interfaces: mouse PcdhγA1 EC1-3 (4zi9), mouse Pcdhα7 EC1-5 (5dzv), mouse Pcdhα4 EC1-4 (5dzw), mouse Pcdhβ6 EC1-4 (5dzx), mouse Pcdhβ8 EC1-4 (5dzy), human PcdhγB3 EC1-4 (5k8r), mouse PcdhγA8 EC1-4 (5szm), mouse PcdhγB7 EC1-4 (5szp), and mouse PcdhγB2 EC1-5 (5t9t) (1, 9–11). This collection of structures yielded a structure-based set of 205 total residues with 1634 residue pairs using the 8 Å cutoff (**Table S4).**

Second, we analyzed our molecular dynamics trajectories to define a simulations-based set of interacting residues, defining an interacting residue pair as a pair of amino acids with non-hydrogen atoms that come within 5 Å of each other in at least 1 frame, 5% or 10% of the simulation frames sampled at 100 ps. From this definition, the interface is defined by a total of 1880 residue pairs originating from a total of 236 interface residues using the 10% cutoff **(Table S4, Figure S9)**. Of the 205 interface residues determined from crystal structures alone, 204 are also in the 236 interface residues from the molecular dynamics simulations, reflecting agreement between these two definitions of clustered Pcdh interface residues. The somewhat expanded simulations-based set likely includes residues that transiently participate in the interface and/or participate in the interface in other isoforms. We thus used this set of 236 interface residues for our sequence-based analyses (**SI Dataset 1**).

**Construction of sequence alignment.** Sequences were found by aligning the PcdhγB3 isoform (Uniprot identifier: PCGDF_HUMAN) against the Uniref database (download date: April 2016) using Jackhmmer with 5 iterations (12). The alignment was filtered to contain only clustered Pcdh sequences, as described in our earlier work (1, 9). The alignment was renumbered according to the mouse Pcdα7 isoform (Uniprot identifier: PCDA7_MOUSE), and gaps were defined relative to this sequence. The alignment was filtered to remove sequences that contain more than 50% gaps and to remove columns that contain more than 50% gaps. The alignment was truncated to contain only domains EC1 through EC4. This alignment has 8560 sequences, with effective number of sequences ($M_{eff}$) of 3300 after down-weighting sequences that are more than 90% identical.

**Iterative pairing algorithm.** Previous work has shown that interacting paralogs from bacteria can be correctly matched by iteratively building a sequence alignment with pairs that have the best $SE_i$ (13, 14). We use a similar approach for the Pcdhs, where we seed an alignment with 1000 randomly paired sequences of EC1-EC2 and EC3-EC4 domains. Each EC1-EC2 is paired with randomly EC3-EC4 from the same species. For speed, we infer couplings using the mean field approximation (15, 16) as implemented in (17). We then assess the $SE_i$ of all possible pairs of sequences within the same species, and keep the top ones for the next iteration. The algorithm was run for 300 iterations, increasing the alignment size by 50 sequences per iteration. Each experiment was repeated for five replicates.

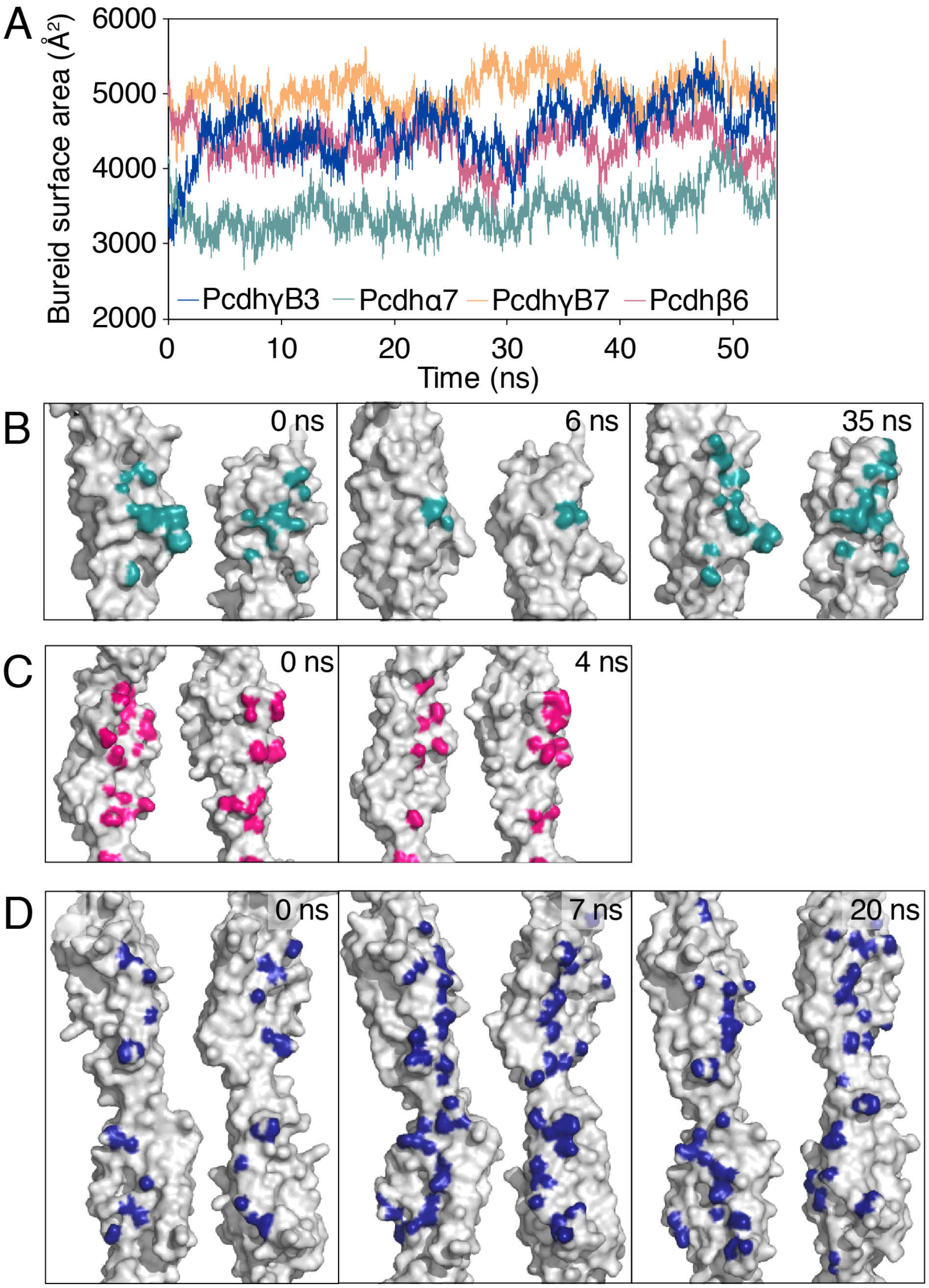


**Figure S1. Clustered protocadherin dimers sample a distribution of conformations**
(A) The overall buried surface areas (BSAs) of the PcdhγB3 (dark blue), PcdhγB7 (orange), Pcdhβ6 (magenta), and Pcdhα7 (teal) complexes are quite large (3400-5000 $Å^2$) and consistent throughout the simulations. (B) The Pcdhα7 EC1/EC4 interaction decreases in BSA (interface residues marked in teal) quickly between 0 and 6 ns, but then increases quickly again at 35 ns, with the interface containing most of the same residues as the initial crystal structure at 0 ns. (C) The Pcdhβ6 EC2/EC3 interaction decreases in BSA from 1400 to 900 $Å^2$ in the first 4 ns of the simulation. (D) The PcdhγB3 EC2:EC3 interaction increases in BSA from 700 to 1400 $Å^2$ in the first 3 ns of the simulation but returns to the lower BSA conformation at 20 ns.

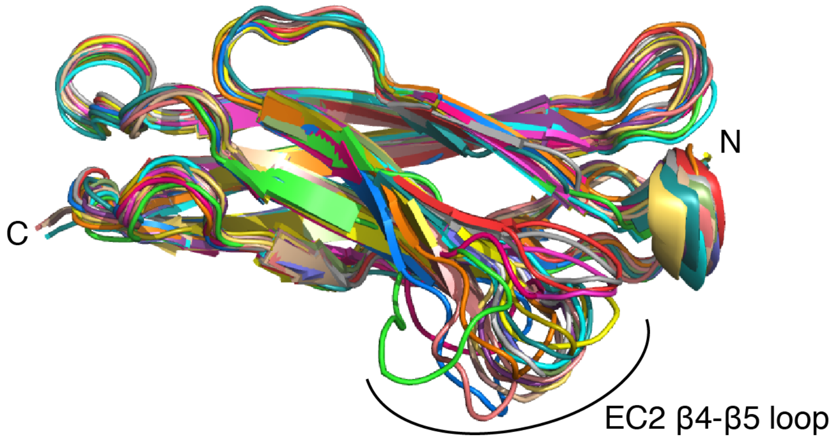


**Figure S2. Conformational diversity of EC2 β4-β5 loop in protocadherin structures**
Superposition of EC2 of the following protocadherin structures shows conformational diversity of the β4-β5 loop:

| | | |
|---|---|---|
| lime green | Pcdhα4 | 5dzw |
| cyan | PcdhγB3 | 5k8r |
| magenta | Pcdhβ1 | 4zpl |
| yellow | Pcdhαc2 | 4zpm |
| light pink | PcdhγC5 | 4zpo |
| white | PcdhγC5 | 4zpp |
| slate blue | PcdhγC5 | 4zpq |
| orange | PcdhγA8 | 4zps |
| teal | Pcdh19 | 5iu9 |
| bright pink | Pcdhα7 | 5dzv |
| light yellow | Pcdhβ6 | 5dzx |
| purple | Pcdhβ8 | 5dzy |
| gray | PcdhγA1 | 5szl |
| sky blue | PcdhγA8 | 5szm |
| gold | PcdhγA9 | 5szn |
| light green | PcdhγC3 | 4zi8 |
| red | PcdhγA1 | 4zi9 |

| | | |
|---|---|---|
| mauve | PcdhγB7 | 5szp |
| light orange | PcdhγB7 | 5szo |
| light red | PcdhγB2 | 5t9t |

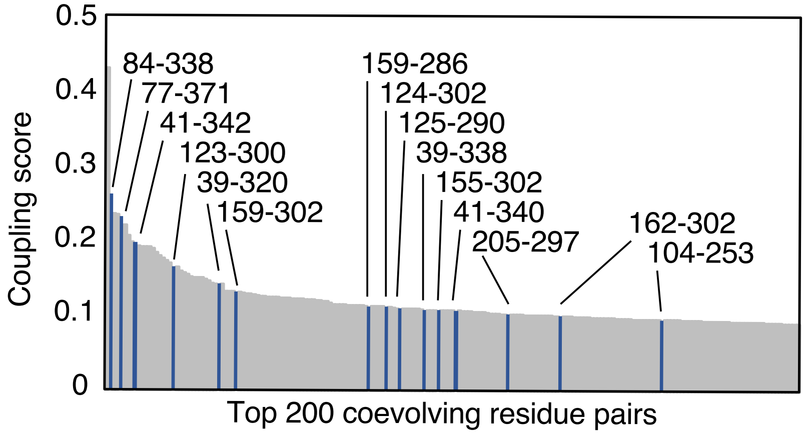


**Figure S3. Rank order of intermolecular coevolving residue pairs.**
Evolutionary coupling scores of the top 200 coevolving residues pairs includes the top 15 intermolecular pairs, highlighted in blue and labeled with corresponding residue numbers in PcdhγB3.

```
        1       10        20        30        40          50        60        70        80        90       99
γB3 EPIRYAIPEELDRGSLVGNLAKDLGFGVGDLPTRNLRVIAEKK--FFTVSPENGNLLVSDRIDREEICGKKSTCVLEFEMVAEKPLNFFHVTVLIQDINDN
α7  GQLHYSVPEEAKHGTFVGRIAQDLGLELTELVPRLFRVASKDRGDLLEVNLQNGILFVNSRIDREELCGRSAECSIHLEVIVDRPLQVFHVEVEVKDINDN
β6  EAIRYSIPEETESGYLVAHLAKDLGFRVGELATRRARIHHRGNKELLQLDVETGNLLLKEKPDREALCGATEPCVLHFQIILENPVQFFQTELQLTDINDH
γB7 QPVRYSIPEELDRGSVVGKLAKDLGLSVLEVSARKLRVSAEKL--HFSVDSESGDLLVKDRIDREQICKGRRKCELQLEAVLENPLNIFHVVVEIEDVNDH

            110       120       130       140       150       160       170       180       190       200     208
γB3 PPTFSQNITELEISELALTGATFALESAQDPDVGVNSLQQYYLSPDPHFSLIQKENLDGSRYPELVLKAPLDREEQPHHHLVLTAVDGGEPSRSCTTQIRVIVADANDN
α7  PPMFPATQKALFILESRLLDSRFPLEGASDADVGSNALLTYRLSTNEHFSLDVPPNHEQVKPLGLVLRKPLDREEAAEIRLLLTATDGGKPELTGTVQLLITVLDVNDN
β6  SPEFPDTEMLLKIQESTQPATVFLLKAAQDSDIGSNAVQNYTVSPNLHFHVVTLSRSDGRKYPELVLDRALDREEQPELTLILTALDGGAPPKSGTTTVRIEVVDINDN
γB7 APQFPKDEINLEISESDSPGARTILESAKDLDIGMNSLSKYQLSPNDYFLLLVKDNPDGSKYPELELQKMLDREAESTHHLMLTAVDGGDPPRTGTTQLRIRVVDANDN

             220       230       240       250        260       270       280       290         300       310
γB3 PPVFTQDMYRVNVAENLPAGSSVLKVMAIDMDEGINAEIIYAFIN-IGKEVRQLFKLDSKTGELTTIGELDFEERDSYTIGVEAKDGG--HHTAYCKVQIDISDENDN
α7  APVFDRSLYTVKLPENVPNGTLVIKVNASDLDEGVNGDVMYSFSSDVSSDIKSKFHMDTVSGEITVIGIIDFEESKAYKIPLEARDKGFPQLPGHCTILVEVVDANDN
β6  APEFVQSLYSVEVPENSPLDALVVTVSARDLDAGIHGNVAYSLFQ-GGGGP-QPFVIDEITGEIRLKGALDFEATSYYTMEIVATDSG--GLSGKCTVAIQVLDVNDN
γB7 RPVFSQDVYRVRLPEDLPPGTTVLRLKAMDQDEGINAEFTYSFLG-VANKAQ--FSLDPITGDIVTRQSLDFEEVEQYTIDVEAKDRG--SLSSQCKVIIEVLDENDN

        320       330       340       350       360       370       380       390       400       410     418
γB3 APEITLASESQHIQEDAELGTAVALIKTHDLDSGFNGEILCQLKGNFPFKIVQDTKNTYRLVTDGALDREQIPEYNVTITATDKGNPPLSSSKTITLHILDVNDN
α7  APQLTVSSLSLPVSEDSQPGRVVTLISVFDRDSGANGQVTCSLTPHIPFKLVSTFKNYYSLVLDSALDRETIANYDVIVTARDGGSPSLWATASVSVEVADVNDN
β6  APKLTISSLTSSIPENAP-EAVVAVFSVSDPDSGDNGRMVCSIQNELPFLLKPTFENYYTLAAEGPLDREIREEYNITIIVSDLGTPRLTTQHTITVQVVDINDN
γB7 RPEIIIITSLSDQISEDSPSGTVVALFKVRDRDSGENAEVMCSLSGNNPFKIHSSSNNYYKLVTDSILDREQTPGYNVTITATDRGKPPLSSSTTITLNVADVNDN
```

**Figure S4. Sequence alignment of clustered protocadherin isoforms**

EC1-4 amino acid sequence and residue numbering of clustered protocadherin isoforms (PcdhγB3, Pcdhα7, Pcdhβ6, and PcdhγB7) on which we performed molecular dynamics simulations.

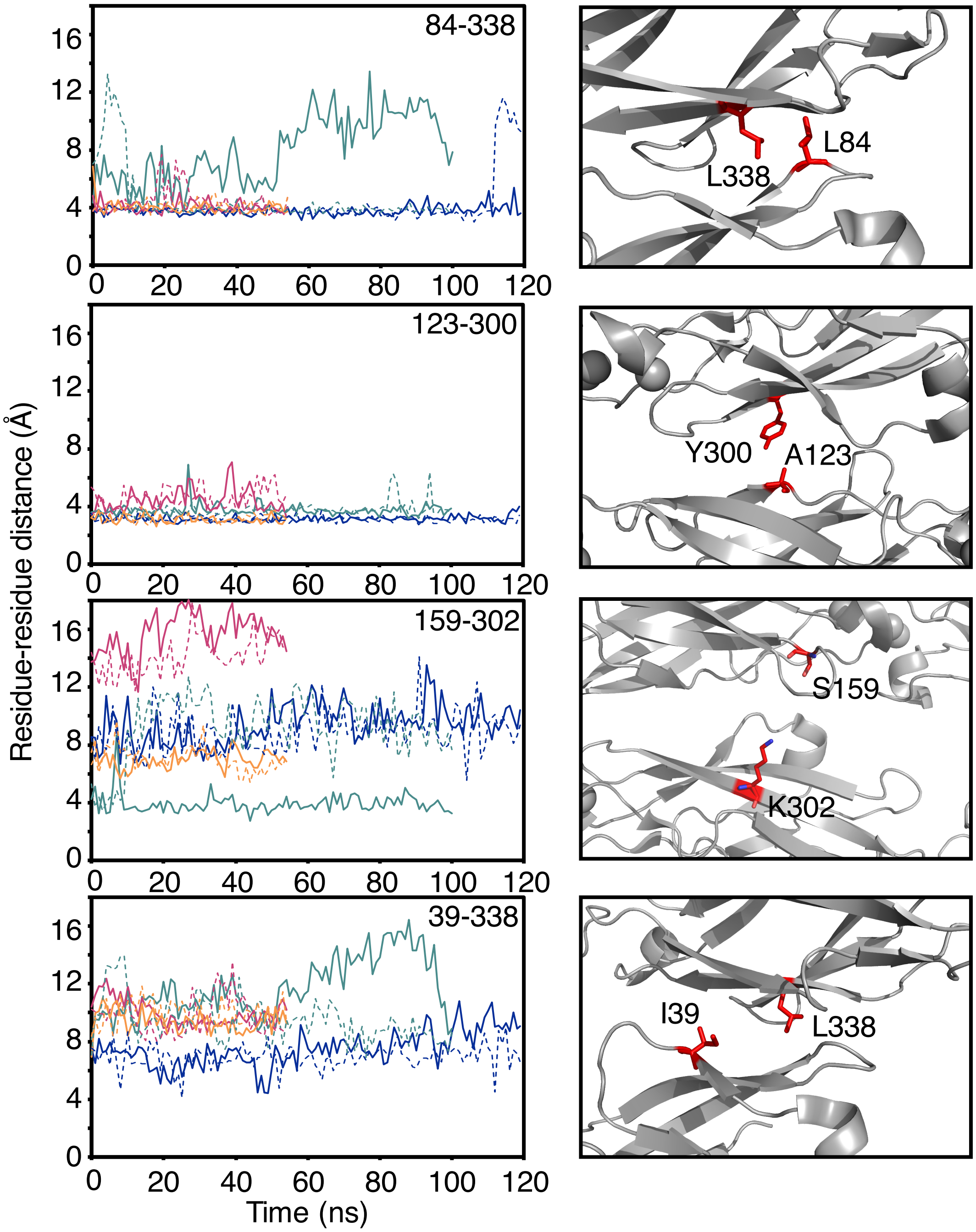


**Figure S5. Diverse residue-residue distance trajectories for coevolving residue pairs** Examples of residue-residue distances plotted over simulation time for some of the top 15 intermolecular coevolving residue pairs. Plots on the left show residue-residue distances for the four simulations (PcdhγB3 (dark blue), PcdhγB7 (orange), Pcdhβ6 (magenta), and Pcdhα7 (teal)) with the two semi-independent pairs from each dimer plotted as solid and dotted lines of the same color. On the right each residue pair in the Pcdh γB3 crystal structure (PDB ID: 5k8r) is illustrated in red stick representation.

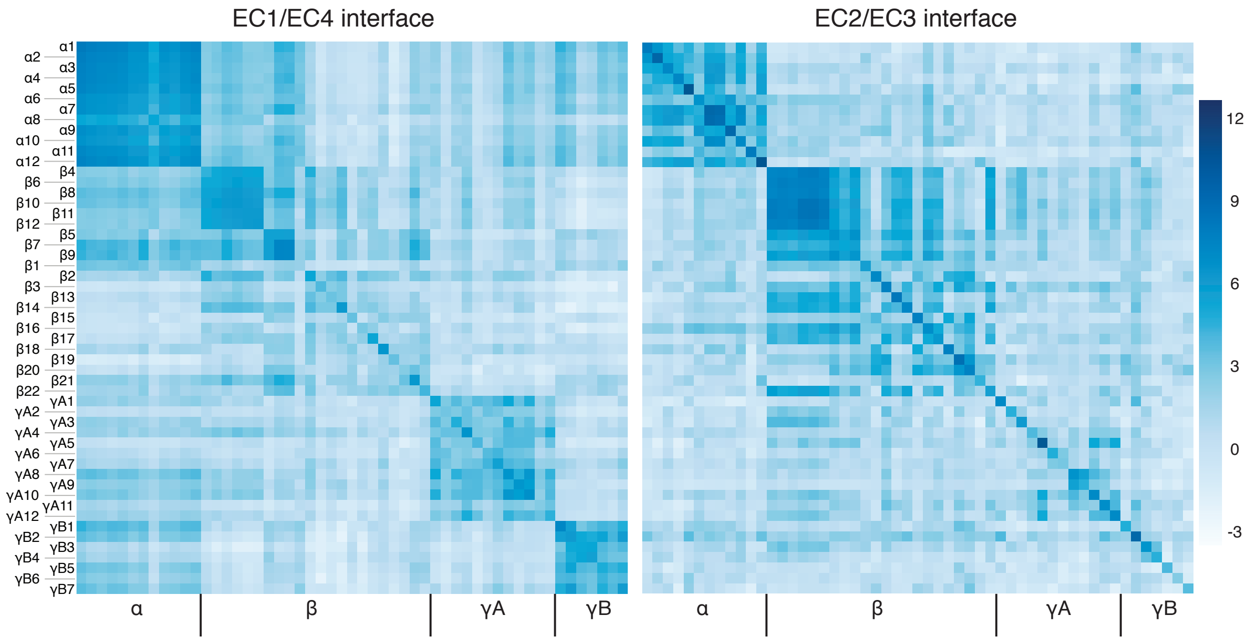


**Figure S6: EC2/EC3 interface $SE_i$ is more distinct between isoforms than EC1/EC4 interface $SE_i$**

$SE_i$ for all possible isoform pairings computed for the EC1/EC4 interface and EC2/EC3 interface. For the α, γA, and γB subfamilies in particular, the EC2/EC3 interface shows high preference for self versus non-self interactions compared to the EC1/EC4 interface.

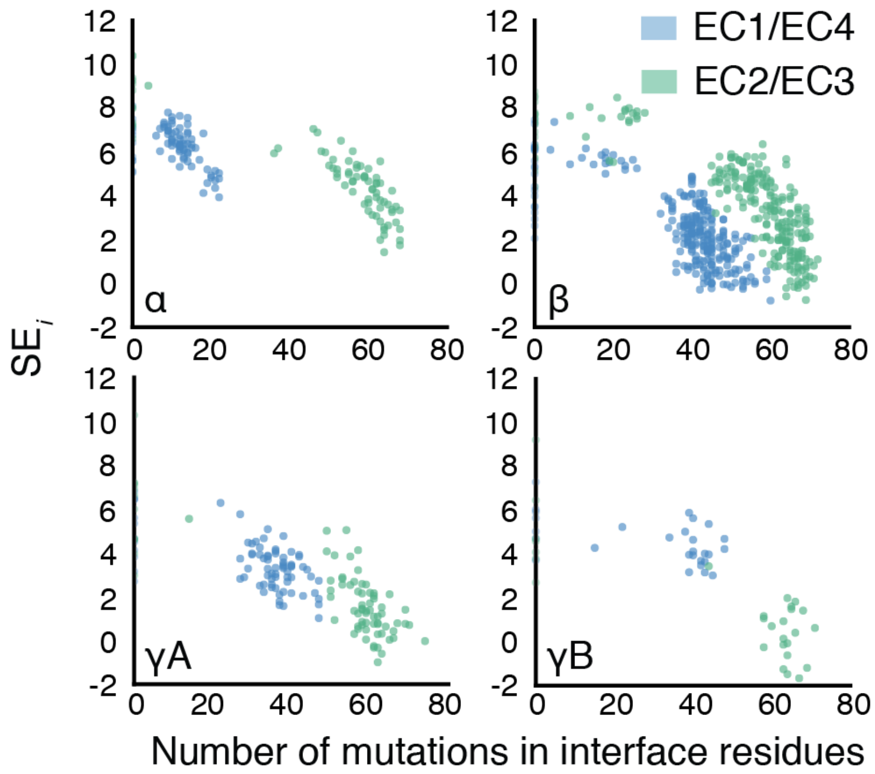


**Figure S7. Statistical energy correlates with number of amino acid substitutions.**
The four plots represent each possible self and non-self interaction of clustered protocadherins within a clustered Pcdh family. The *x*-axis shows the number of mutations that separate each non-self pair, and the *y*-axis shows the statistical energy of interaction ($SE_i$) of the corresponding interaction interface. In general, the more mutations separate the two Pcdh pairs, the lower the predicted statistical energy.

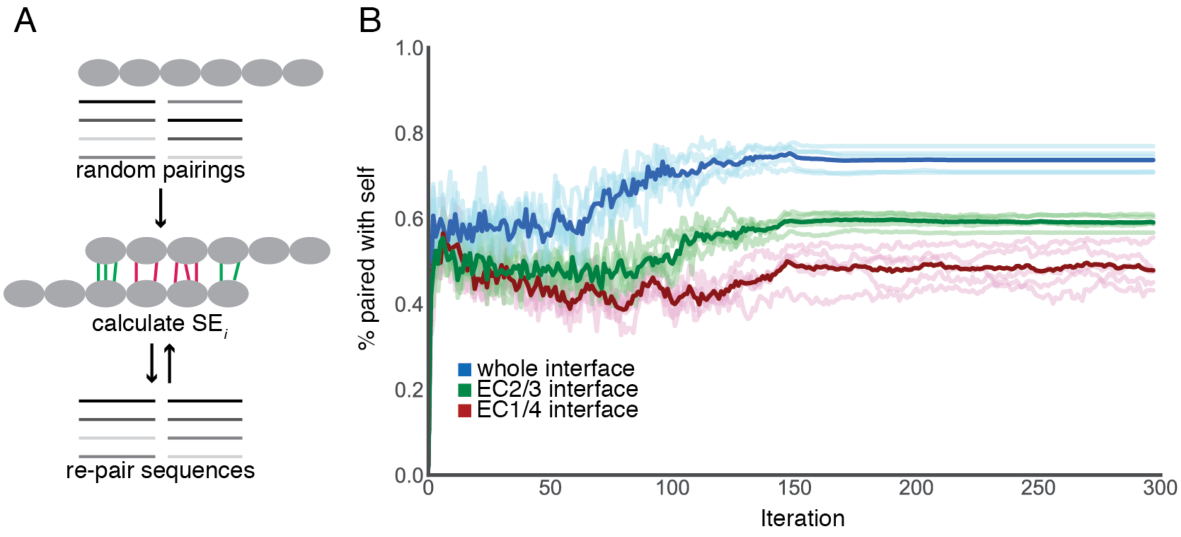


**Figure S8. Iterative pairing algorithm establishes that EC2/EC3 contributes more to specificity**

(A) Schematic of the iterative pairing algorithm. After randomly pairing starting sequences, the parameters of the evolutionary couplings model are inferred and used to update the pairings. Each cycle of inference and update is one iteration. (B) Results of five replicates where the evolutionary couplings from the whole interface, just the EC2/EC3 interface, or just the EC1/EC4 interface are used to update the alignments. Light lines indicate individual runs, dark runs are the mean of the replicates.

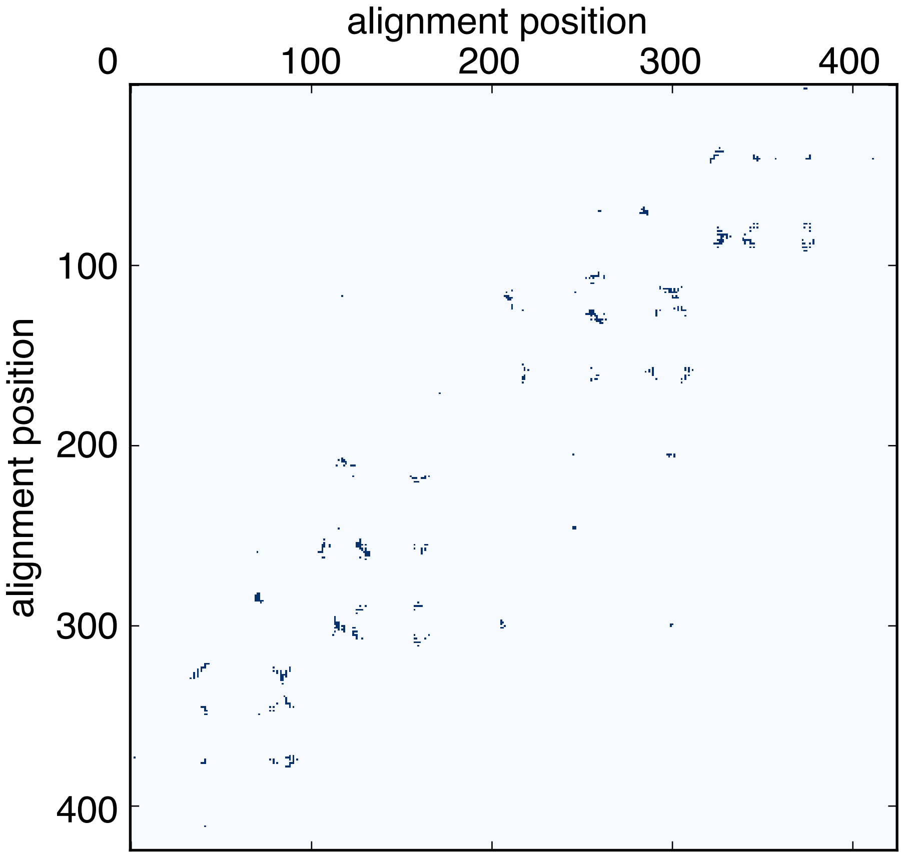


**Figure S9. Interface contact map from Pcdh simulations**
Dimer interface contact map, where each dot represents two residues that come within 5 Å of each other (distance measured at the closest pair of non-hydrogen atoms) for at least 10% of the simulations of human PcdhγB3 EC1-4, mouse PcdhγB7 EC1-4, mouse Pcdhα7 EC1-5, or mouse Pcdhβ6 EC1-4. The resulting 1880 pairs arise from a total of 236 residues which comprise the interface residues used to generate the statistical interaction energy model.
<insert page break then Fig. S1 here>

**Table S1. Overview of MD simulations**

| Label | PDB | $t_{sim}$ (ns) | Size (# atoms) | Size ($nm^3$) |
|---|---|---|---|---|
| Mouse PCDHα7 EC1- EC5 | 5dzv | 100.0 | 500,917 | 52.4 x 9.94 x 9.45 |
| Mouse PCDHβ6 EC1-EC4 | 5dzx | 60.6 | 394,722 | 41.5 x 9.86 x 9.48 |
| Human PCDHγB3 EC1-EC4 | 5k8r | 120.0 | 335,584 | 48.9 x 8.21 x 8.21 |
| Mouse PCDHγB7 EC1-EC4 | 5szp | 61.2 | 308,093 | 43.5 x 8.84 x 78.8 |

**Table S2. BSA of individual EC interactions in MD simulations**

| Isoform | Interacting pair | BSA (Å$^2$) | Overall BSA (Å$^2$) |
|---|---|---|---|
| PcdhγB3 | EC1/EC4 | 900±100 | 4600±300 |
| | EC2/EC3 | 1200±200 | |
| | EC3/EC2 | 1000±200 | |
| | EC4/EC1 | 900±100 | |
| Pcdhα7 | EC1/EC4 | 700±200 | 3400±300 |
| | EC2/EC3 | 900±100 | |
| | EC3/EC2 | 600±100 | |
| | EC4/EC1 | 1000±300 | |
| Pcdhβ6 | EC1/EC4 | 1000±100 | 4300±200 |
| | EC2/EC3 | 1000±100 | |
| | EC3/EC2 | 1100±100 | |
| | EC4/EC1 | 900±100 | |
| PcdhγB7 | EC1/EC4 | 1000±100 | 5100±200 |
| | EC2/EC3 | 1300±100 | |
| | EC3/EC2 | 1300±100 | |
| | EC4/EC1 | 900±100 | |

**Table S3. Data statistics for low HEPES and HEPES-free PcdhγB3 EC1-4 structures**

| Protein | PcdhγB3 EC1-4 no HEPES | PcdhγB3 EC1-4 less HEPES |
|---|---|---|
| PDB ID | 6meq | 6mer |
| SBGridDB ID | 602 | 603 |
| Data Collection | | |
| Beam source | APS 24-ID-C | APS 24-ID-C |
| Wavelength (Å) | 1.07 | 0.98 |
| Space group | $C222_1$ | $C222_1$ |
| Unit cell (a, b, c; Å) | 128.39, 161.77, 52.16 | 126.81, 162.91, 52.86 |
| Unit cell (α, β, γ) | 90, 90, 90 | 90, 90, 90 |
| Resolution (Å) | 28.61-3.0 (3.107-3.0) | 46.74-2.9 (3.004-2.9) |
| Total reflections | 39257 (3576) | 61580 (2636) |
| Unique reflections | 10828 (1046) | 11833 (819) |
| Multiplicity | 3.6 (3.4) | 5.2 (3.2) |
| Completeness (%) | 95.75 (93.97) | 94.21 (67.19) |
| Mean I/σ(I) | 9.36 (1.60) | 10.02 (1.45) |
| Wilson B-factor | 82.12 | 75.67 |
| $R_{merge}$ | 0.147 (0.987) | 0.113 (0.680) |
| $R_{meas}$ | 0.167 (1.15) | 0.125 (0.793) |
| $CC_{1/2}$ | 0.988 (0.51) | 0.996 (0.617) |
| CC* | 0.997 (0.822) | 0.999 (0.874) |
| Refinement | | |
| Refinement resolution range | 28.61-3.0 (3.107-3.0) | 46.74-2.9 (3.004-2.9) |
| Reflections used in refinement | 10825 (1045) | 11831 (819) |
| Reflections used for R-free | 1089 (104) | 1181 (78) |
| $R_{work}$ | 0.225 (0.336) | 0.224 (0.377) |
| $R_{free}$ | 0.272 (0.390) | 0.271 (0.446) |
| $CC_{work}$ | 0.952 (0.638) | 0.940 (0.600) |
| $CC_{free}$ | 0.911 (0.449) | 0.877 (0.279) |
| Number of non-hydrogen atoms | 3326 | 3330 |
| Macromolecules | 3219 | 3229 |
| Ligands ($Ca^{2+}$) | 9 | 9 |
| Waters | 98 | 92 |
| Protein residues | 414 | 416 |
| RMS Bonds (Å) | 0.003 | 0.003 |
| RMS Angles (°) | 0.90 | 0.88 |
| Clashscore | 3.45 | 3.12 |
| Average B-factor | 92.38 | 92.28 |
| Macromolecules | 93.47 | 93.25 |
| Ligands | 74.16 | 75.77 |
| Solvent | 58.32 | 59.78 |
| Ramachandran plot regions | | |
| Favored (%) | 96.60 | 96.38 |
| Allowed (%) | 3.40 | 3.62 |
| Outliers (%) | 0 | 0 |
| Rotamer outliers (%) | 0.28 | 1.93 |

**Table S4. Interface residues from Pcdh structures and simulations**

| Grouping | number of pairs | number of positions |
|---|---|---|
| crystal 5 Å* | 542 | 141 |
| crystal 8 Å* | 1634 | 205 |
| MD 10%‡ | 1880 | 236 |
| MD 5%‡ | 2096 | 243 |
| MD 0%‡ | 3042 | 276 |

* Residue pairs that are within 5 or 8 Å of each other in crystal structures

‡ Residue pairs that are within 5 Å of each other for at least 0%, 5% or 10% of the simulation frames

**Additional dataset S1 (separate file)**

List of interface residues from molecular dynamics simulations based on PcdhγB3 numbering

**Additional dataset S2 (separate file)**

List of interface residue pairs used in statistical energy of interaction model based on PcdhγB3 numbering

**Additional dataset S3 (separate file)**

Alignment of mouse clustered Pcdh isoforms
